# ECM cross-linking regulates invadopodia dynamics

**DOI:** 10.1101/198580

**Authors:** Kamyar Esmaeili, Aviv Bergman, Bojana Gligorijevic

## Abstract

InvadopodiInvadopodia are membrane protrusions dynamically assembled by invasive cancer cells in contact with extracellular matrix (ECM). Invadopodia are enriched for the structural proteins actin and cortactin, as well as metalloproteases such as MT1-MMP, whose function is to degrade the surrounding ECM. During metastasis, invadopodia are necessary for cancer cell intravasation and extravasation. While signaling pathways involved in the assembly and function of invadopodia are well studied, few studies address invadopodia dynamics and how the cell-ECM interactions contribute to cell invasion. Using iterative analysis based on time-lapse microscopy and mathematical modeling of invasive cancer cells, we found that cells oscillate between invadopodia presence and cell stasis, termed Invadopodia state and invadopodia absence during cell translocation, termed Migration state. Our data suggests that β1-integrin-ECM binding and ECM cross-linking control the duration of each of the two states. By changing the concentration of cross-linkers in 2D and 3D cultures, we generate ECM where 0-0.92 of total lysine residues are cross-linked. Using ECM with a range of cross-linking degrees we demonstrate that the dynamics of invadopodia-related functions have a biphasic relationship to ECM cross-linking. At intermediate levels of ECM cross-linking (0.39), cells exhibit rapid invadopodia protrusion-retraction cycles and rapid calcium spikes, which lead to more frequent MT1-MMP delivery, causing maximal invadopodia-mediated ECM degradation. In contrast, both extremely high or low levels of cross-linking lead to slower invadopodia-related dynamics and lower ECM degradation. Additionally, β1-integrin inhibition modifies dynamics of invadopodia-related functions, as well as the length of time cells spend in either of the states. Collectively, these data suggest that β1-integrin-ECM binding non-linearly translates small physical differences in extracellular environment to differences in the dynamics of cancer cell behaviors. Understanding conditions under which invadopodia can be reduced by subtle environment-targeting treatments may lead to combination therapies for preventing metastatic spread.

## Introduction

Invadopodia (1) are dynamic membrane protrusions involved in the invasive motility of cancer cells. Their function is to degrade the surrounding ECM. In tumors *in vivo*, they are necessary for cancer cell intravasation (2) and extravasation (3) from the blood vessels. *In vitro*, invasive cancer cells assemble invadopodia when grown on various ECM components and in the presence of growth factors. Invadopodia have appearance of puncta rich in actin, actin-regulatory proteins (e.g. cortactin), tyrosine kinases and proteases (eg. MT1-MMP). While a large number of the invadopodial components are found in other motility-related structures such as focal adhesions and lamelipodia, the unique feature of invadopodia is the high ECM-degrading activity.

Assembly and maturation of an individual invadopodium has been well studied (4, 5) using 2D *in vitro* assays. Invadopodium precursors, consisting of cortactin, cofilin and N-WASP, are first assembled (6). The structure is then stabilized via focal adhesion proteins such as β1-integrin by binding the structure tip to the ECM (7, 8). During invadopodia maturation, cortactin phosphorylation leads to continuous actin polymerization and MT1-MMP recruitment to the tip of invadopodia via late endosomes (9). Transmembrane MT1-MMP assures focalized ECM degradation of the surrounding ECM and invadopodia becomes mature and functional (at ˜50 minutes) (6). During ECM degrading activity, mature invadopodia exhibits SOCE (Store-operated calcium entry)-dependent calcium spikes, which are necessary for MT1-MMP recycling to the plasma membrane (10). Mature invadopodia also exert physical force on ECM, by means of protrusion-retraction cycles, during which actin filaments polymerize (protrusion) and depolymerize (retraction) repeatedly (11). Physical and proteolytic ECM remodeling by invadopodia are coordinated via cortactin (de)phosphorylation and positive feedback from MT1-MMP (12, 13).

The assembly of invadopodia, as well as level of ECM degradation were shown to be linked to the ECM properties such as rigidity, density and cross-linking (14–16). This points to the essential role in adhesion-based signaling in the invadopodia progression. A recent study of β1-integrin role in invadopodia strengthened this link, demonstrating that β1-integrin is localized to invadopodia and directly links the structure to ECM (7). While elimination of β1-integrin does not inhibit invadopodia precursor assembly, it disables invadopodia maturation and ECM-degrading function.

Recent studies of invadopodia in 3D (13, 17, 18) and *in vivo* (2, 19) demonstrated that invadopodia are present at the front of the cells migrating in MMP-dependent manner. Depending on the ECM parameters including density, fiber size, stiffness and/or cross-linking, such migration can result in a broad range of speeds (6-30 μm/hour for breast carcinoma) (13, 20, 21). *In vivo*, presence of invadopodia was also shown to be highly dependent on ECM cross-linking (2). While the studies of 3D and *in vivo* invadopodia established the link between invadopodia and cell movement, it is not known if cells which assemble invadopodia can translocate the cell body simultaneously or they move and assemble invadopodia in a sequential manner.

We have used iterative cycles of time-lapse microscopy and mathematical modeling towards addressing the role of invadopodia dynamics and ECM cross-linking in invasive cell motility. We found that the dynamics of invadopodia assembly and function have a non-monotonic (biphasic) relationship to an increase in ECM cross-linking. Furthermore, we show that the presence of invadopodia and the migration of cells are negatively correlated, with level of active ß1-integrin controlling the duration of each of the two states. We demonstrate that partial blocking of β1-integrin increases the duration of migration and shortens the period of active ECM degradation by inhibiting invadopodia activity. Taken together, our results suggest that invadopodia-driven, MMP-dependent motility consists of oscillations between a) sessile invadopodia assembly leading to ECM degradation and b) migration. Modulation of β1-integrin activity or ECM cross-linking can be used to reduce or eliminate invadopodia, which are necessary for intravasation and metastasis (2).

## Results

### Breast cancer cells switch from migration to invadopodia-mediated ECM degradation

To determine whether cell migration occurs simultaneously or sequentially with invadopodia assembly, we acquired time-lapse recordings of >100 cells in each of the two breast carcinoma cell lines known to spontaneously assemble invadopodia (MTLn3 and MDA-MB-231). Cells were engineered to stably express fluorescent invadopodia markers (Cerulean-cortactin or Ceruelan-LifeAct) and cultured on fluorescent Alexa 488-gelatin (**Movie S1**, **S2**). Recordings were scored by tracking centroid path over time (**Figure 1A-C**, path) as well as quantifying number of invadopodia (**Figure 1C**, cyan punctae).

**Figure 1.**
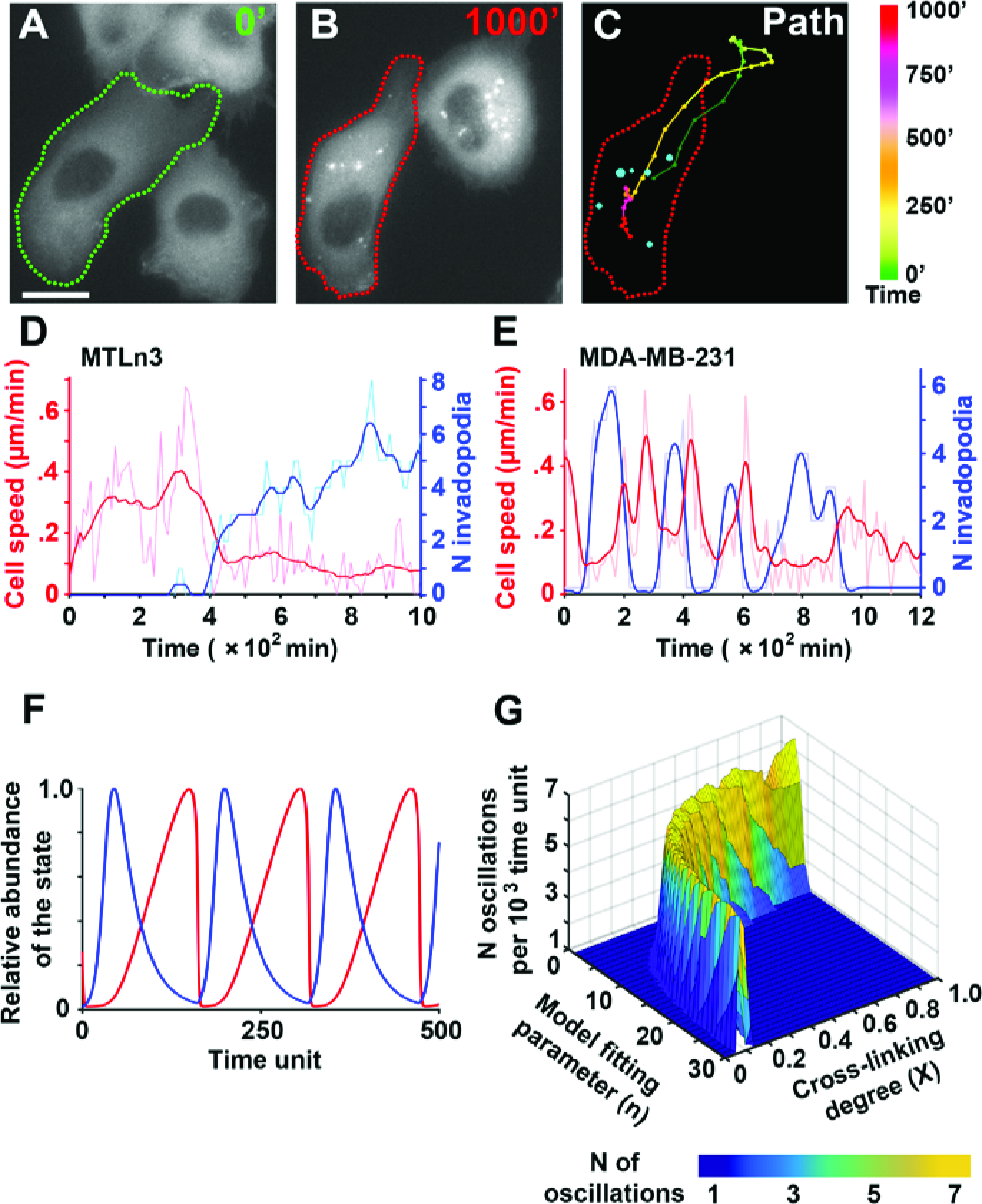
Oscillations between Migration and Invadopodia states in cancer cells and the role of ECM cross-linking: **A.** The micrograph showing Cerulean-cortactin-MTLn3 cell at time 0 of the **Movie S1**. Cell border is highlighted with the green dotted line. Scale bar 10 um; **B.** The same cell at 1000 minutes highlighted in red; **C.** Summary of A and B, with color-coded path 0-1000 minutes and cyan punctae representing invadopodia at 1000 minutes. Color-code legend is shown on the right. **D & E.** Quantification of **Movie S1** (Cerulean-cortactin-MTLn3) and **Movie S2** (Cerulean-LifeAct-MDA-MB-231) respectively. Cell speed in μm/min (red lines) and number of invadopodia (blue lines) were recorded over time in representative cells. Raw data (shaded lines), smoothened data (bright lines); **F.** Simulation of cancer cell oscillations between invadopodia (blue, relative abundance I/I_max_ vs time) and migration (red, relative abundance M/M_max_ vs time). **G.** Results of the model: Number of invadopodia-migration oscillations shows a biphasic trend to linear changes in ECM cross-linking and model fitting parameter n.

In the beginning of **Movie S1**, the tracked cell migrates relatively fast (>0.3 μm/min or >18 μm/h), (**Figure 1D**, red trace). While cell assembles several short-living (<10 min lifetime) invadopodia precursors, characterized by cortactin-enriched punctae, none of them result in any detectable ECM degradation. Starting at 380 minutes, speed of the cell decreases (<0.2 μm/min) and the cell assembles mature invadopodia, characterized by cortactin-enriched punctae with >50 min lifetime colocalized with ECM degradation (**Movie S1**, red circles and **Figure 1D**, blue trace). Similar observations were next confirmed in MDA-MB-231 cells (**Movie S2**), where several cycles of switching between migration and mature invadopodia were observed (**Figure 1E**).

Based on these observations, we defined two oscillating cell states: Migration state, where cells exhibit speeds >0.3 μm/min and no ECM degradation, and Invadopodia state, where cells exhibit punctae rich in cortactin (or actin) colocalized with ECM degradation and migrate at speeds <0.2 μm/min.

Our previous work in vivo suggested that the balance between migration and invadopodia is determined by ECM cross-linking (2). Moreover, ECM cross-linking has been demonstrated to enhance integrin signaling (22) and increase the levels of ECM degradation by invadopodia (15). Hence, we hypothesized that the frequency of switching between Invadopodia state and Migration state will be controlled by ECM cross-linking and integrin-driven ECM-cell interactions (20–25). To develop a generalizable prediction, we developed a phenomenological, non-dimensional mathematical model (see **Supplemental Information**), based on the observed state oscillations and prior studies on ECM-cell interactions in 2D, 3D and *in vivo* environments. The dynamic variables used in the model were the concentration of ECM interacting with the cell (represented by CECM), abundance of Invadopodia state (represented by I) and abundance of Migration state (represented by M). Cell-ECM interactions were assumed to be constant for specific ECM conditions and addressed by the adhesion constant Ka. To address the dynamic properties of the migration/degradation oscillations, we varied the degree of ECM cross-linking (expressed as X, the ratio of cross-linked lysine residues to the total number of lysine residues). We additionally introduced model fitting parameter (n), which is used to fit the model to different ECM dimensionalities. Parameter n was further experimentally determined to be 6 in 2D assays, and 7 in 3D conditions (**Figure S1**).

**Figure 1F** shows one run of the model simulation for a cell oscillating between invadopodia (blue line) and migration (red line). The dynamics of the transition between Invadopodia and Migration states observed at a single cell level (**Figure 1D, E**) is recapitulated and extended in the repeated cycles of migration/invadopodia that model demonstrates. **Figure 1G** summarizes the model simulations for varying X, n and oscillation frequencies. The model suggests that an increase in ECM cross-linking will enable a biphasic change in the frequency of migration/invadopodia switches in cells. Such a prediction implies that at an intermediate cross-linking X, the number of switches from migration to degradation and vice versa will reach a maximum (**Figure 1G**, yellow regions), which will in turn result in the maximal number of degraded punctae.

### Invadopodium maturation and ECM degradation biphasically conform to ECM cross-linking in 2D and 3D

To test if the prediction of our model is supported in experimental invadopodia assays, we measured invadopodia assembly and function in ECM with increasing cross-linking degrees. Breast carcinoma MTLn3 cells were plated on a thin fluorescent gelatin layer cross-linked from 0 to 0.92 cross-linking degree (**Figure S2A**), using the chemical cross-linker glutaraldehyde (GTA). At 18 hours after plating, the number of invadopodia precursors, mature invadopodia and ECM degradation were measured (**Figure 2A-D**). The number of invadopodia precursors reached a plateau at 0.39 cross-linking degree (**Figure 2B**), while the number of mature invadopodia showed a strong biphasic trend with the increase in cross-linking (**Figures 2C, D**). The peak of the biphasic trend is at the intermediate cross-linking level (0.39) for both the total number of spots degraded over 18 hours (**Figure 2C**) and for mature invadopodia present at the time of cell fixation (**Figure 2D**). Moreover, the portion of invadopodia precursors that mature and degrade ECM, reported as invadopodia stability ratio, is at its maximum at the cross-linking degree of 0.39 (**Figure 2E**). Trend in increase of total area of degradation further confirms that the increase in the cross-linking degree has a biphasic relationship to invadopodia maturation and degradation. Our results were also confirmed using two additional human breast carcinoma cell lines, MDAMB-231 and Hs-578T (**Figure 2F**). All three cell lines presented biphasic distributions of invadopodia-based degradation with increased cross-linking, with the peak at the intermediate cross-linking (X = 0.39). In conclusion, as predicted in our model, the experimental data show a biphasic trend in invadopodia degradation with increased cross-linking ratio.

**Figure 2.**
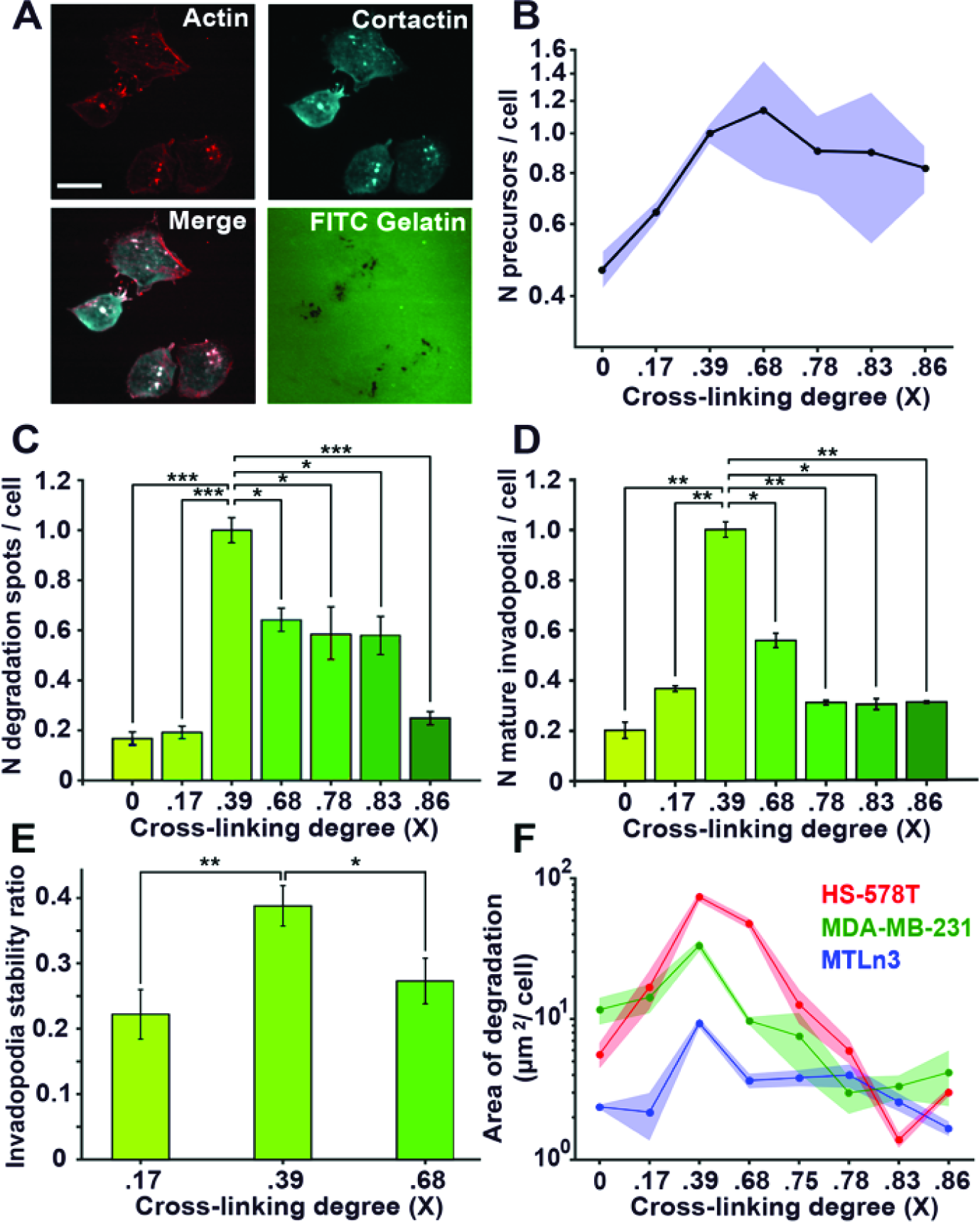
ECM cross-linking biphasically regulates invadopodia maturation and degradation: **A.** Representative fluorescence micrographs showing colocalization of actin (red) and cortactin (cyan) in invadopodia of MTLn3 cells. Lower right panel shows degradation of gelatin at 18 hours. Scale bar 10 μm; **B-D.** Relative number of invadopodia precursors (**B**), mature invadopodia (**C**) and number of degradation spots (**D**) at different cross-linker (GTA) concentrations in MTLn3 cells. In (**B-D**), bars represent >300 cells in 3 separate experiments per condition; data are normalized to 0.05 % GTA. (**E**) Invadopodia stability ratio at three concentrations of cross-linker; bars represent >200 invadopodia in 3 separate experiments per condition; SEM errors are shown. (**F**) Area of degradation measured at 18 hours in three breast carcinoma cell lines: MTLn3 (blue), MDA-MB-231 (green) and Hs578 (red). Experiments are performed in triplicate and circles represent average degradation area per cell in >10 fields of view at each repeat. Lines represent mean values and shaded areas represent 95% confidence intervals. In **B** and **F**, vertical axes are in logarithmic scale. All p values report comparison to 0.39 cross-linking ratio condition. * p<0.05; **p<0.01, *** p<0.001.

Results of the 2D invadopodia assay were then validated in 3D invadopodia assay, using FITC-DQ-collagen cross-linked using transglutaminase II from a 0 to 0.87 crosslinking degree (**Figure S2B**) (2, 13, 26, 27). Colocalization of the FITC-DQ collagen I signal with cortactin, actin, and MT1-MMP confirmed that the 3D collagen degradation was due to invadopodia activity (**Figure 3A**). Interestingly, invadopodia-mediated degradation also followed a biphasic trend in a 3D environment, with maximum degradation present at an intermediate level of crosslinking (0.36) (**Figure 3B**).

**Figure 3.**
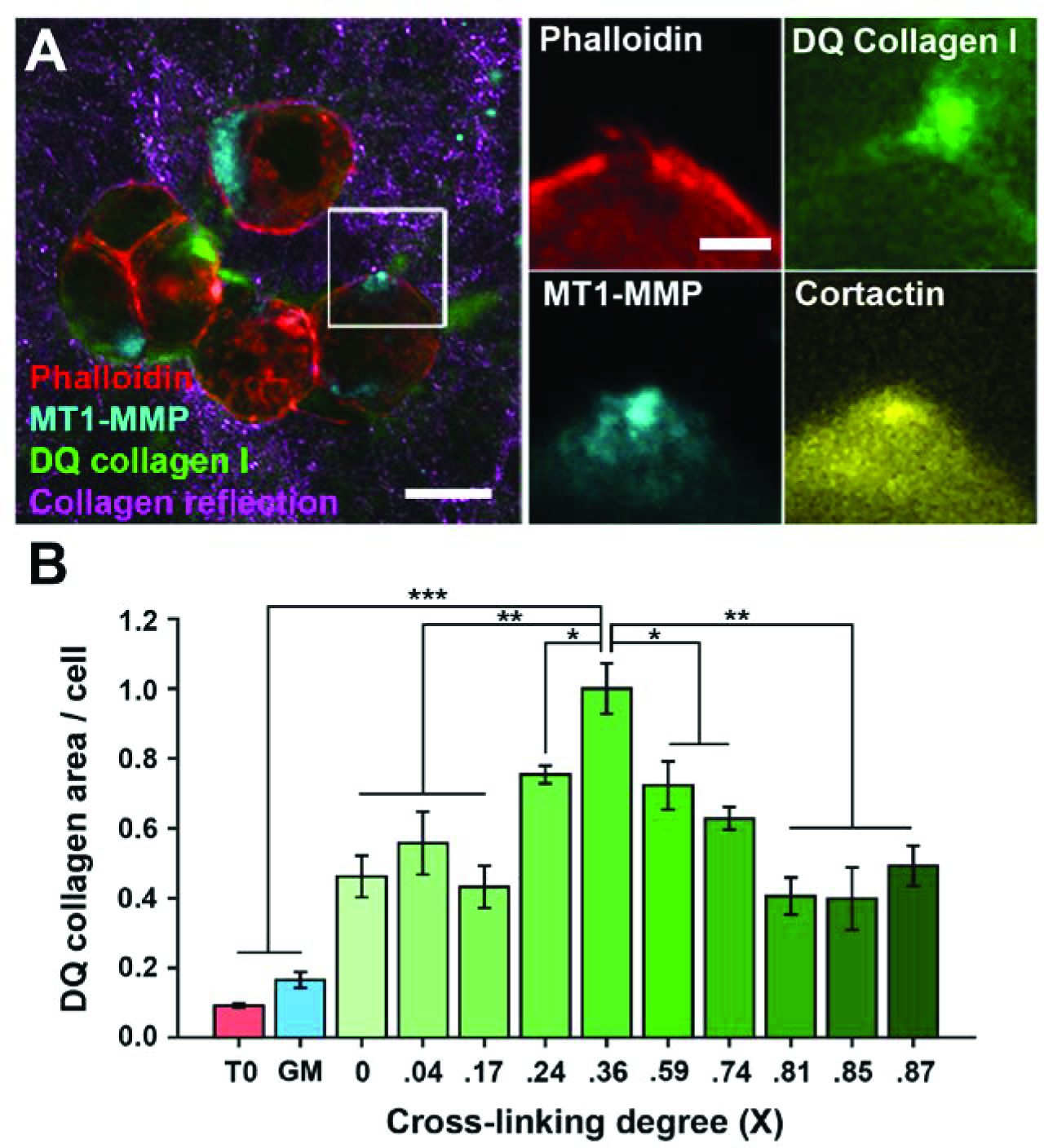
Collagen I cross-linking regulates ECM remodeling and degradation in 3D: **A.** Representative fluorescence images depicting MTLn3 cells in 3D collagen I matrix. After 42 hours, cells are fixed and immunostained for actin (red), cortactin (yellow), and MT1-MMP (cyan). Collagen fibers were detected with reflected confocal microscopy (magenta) and collagen degradation was measured using FITC-DQ collagen I (green). **B.** FITC-DQ collagen-positive areas in 3D cultures. Green bars present FITC-DQ-collagen-positive areas measured at 42h post-embedding into collagen with different cross-linking degrees. Red bar (T0) represents 0h; blue bar (GM) represents culture after 42h treatment with GM6001, both at cross-linking degree of 0.36. Experiments are performed in triplicate and bars represent average area per cell in >10 fields of view at each repeat. All p values report comparison to the condition with 0.36 cross-linking ratio. * P<0.05; ** P<0.01, *** P<0.001.

Furthermore, measurements of collagen gel pore size and storage modulus revealed monotonic variation of these variables by increasing ECM cross-linking (**Figure S2C, D**). Biphasic variations of invadopodia-mediated ECM degradation as a response to monotonic changes of ECM cross-linking, pore size and storage modulus pretermits passive regulation of invadopodia activity by ECM stiffening and suggests that invadopodia are actively responding to these changes at the molecular level.

Collectively, our model and experimental, end-point measurements showed that the intermediate cross-linking degree results in the increase of total number of invadopodia maturing and degrading ECM over time. We hypothesized that this can be a result of faster dynamics of invadopodia activities, such as faster protrusive cycles and/or more frequent delivery of MT1-MMP.

### Frequency of invadopodium protrusion-retraction cycles changes biphasically with increase in ECM cross-linking

To assess the effect of ECM cross-linking on invadopodia protrusion-retraction cycles, fluorescence intensity of cortactin punctae (**Movie S3**) was monitored in MTLn3 cells at various gelatin cross-linking degrees (**Figure 4)**. Cortactin signal oscillations at invadopodia, reflecting cycles of protrusion-retraction, were filtered (**Figure 4B**) and analyzed by autocorrelation (**Figure 4C**). Autocorrelation algorithm correlates the signal of the cortactin oscillations with delayed copies of itself in order to find the periodicity of a repeating pattern within the signal. Results show average frequencies of protrusion-retraction cycles of approximately 2.5 mHz at low or high (0.18 or 0.69) cross-linking degrees and significantly higher frequency of 3.08 mHz at intermediate cross-linking (**Figure 4D**). This reflects significantly faster protrusion-retraction cycles and suggests that the dynamics of invadopodia protrusive cycles is in direct relationship to ECM cross-linking (**Figure 4D**). Furthermore, inhibiting F-actin polymerization by 4 μM Cytochalasin D resulted in total abrogation of cortactin oscillations (**Movie S4** and **Figure 4E, F**), confirming that measured oscillations in cortactin signal reflect cycles of invadopodia protrusions and retractions. Combined with results shown in Figures 2 and 3, this data indicates that increased degradation at intermediate ECM cross-linking may be the consequence of faster protrusive cycles.

**Figure 4.**
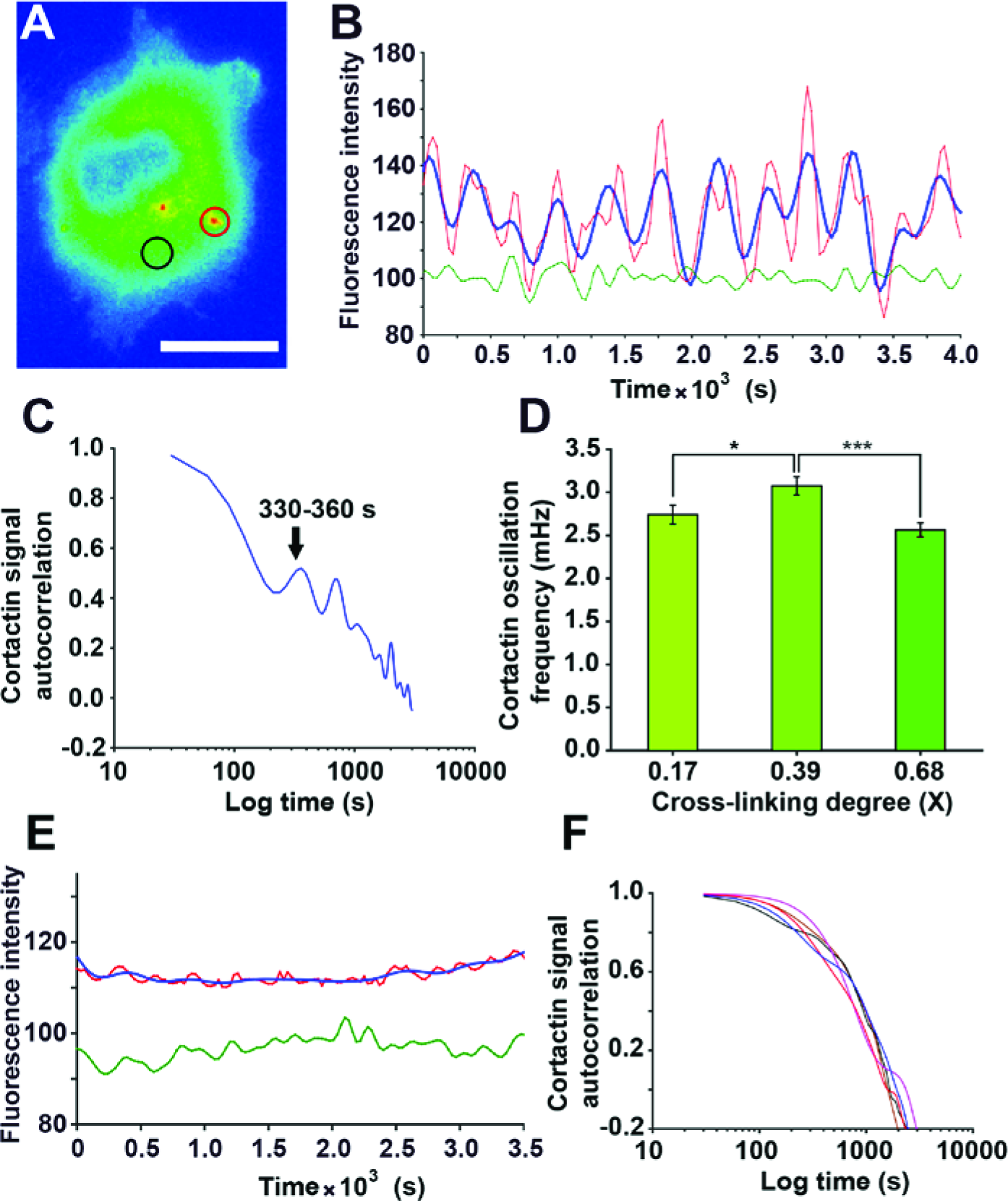
Frequency of cortactin oscillations is biphasically regulated by ECM cross-linking. **A.** Representative pseudocolored image of Cerulean-cortactin-MTLn3 cell. Invadopodium is circled in red, while same size of area in the cytoplasm is circled in black. Signal inside cytoplasmic circle is used as a filter for cortactin signal in the invadopodium**; B.** Representative traces of cortactin signal oscillations (red line), cytoplasmic signal representing the noise (green line) and filtered cortactin signal (blue line) corresponding to areas within the circles in (**A**); **C**. Autocorrelation analysis of the filtered cortactin signal showing the periodicity of the signal. **D.** Bars show the cortactin oscillation frequency at different ECM cross-linking and represent >10 cells per experiment and 3 separate experiments per condition; **E**. Cortactin fluorescence over time measured in a cell treated with F-actin inhibitor Cytochalasin D (**Movie S4**). Fluorescence was measured in the invadopodia precursor (red line, prior to filtering; blue line, post-filtering) and in the cytoplasm (green line). **F**. Autocorrelation of cortactin signals in 5 cells with invadopodia precursors shows cortactin oscillations are absent when F-actin polymerization is inhibited. Pairwise comparisons were done with 0.39 cross-linking ratio condition. Errors are shown as SEM. * P<0.05; ** P<0.01.

Next, we tested if the increase in ECM cross-linking affects proteolytic degradation of ECM by increased dynamics of calcium spikes and consequent MT1-MMP recycling to the plasma membrane (10).

### Calcium spiking frequency biphasically changes with increase in ECM cross-linking

SOCE-dependent calcium spikes were recently shown to be essential both for precursor assembly via Src activation, as well as MT1-MMP recycling to the plasma membrane during ECM degradation (10). As the calcium channels are commonly mechanosensitive (28–30) and calcium spiking can be influenced by ECM (31), we hypothesized that the frequency of calcium spikes will reach a peak value at cross-linking degree of 0.39, in coordination with protrusion-retraction dynamics. To test this, we monitored calcium spikes in MTLn3 cells with invadopodia, recording Fluo-4-AM signal at different cross-linking degrees (**Movie S5**; **Figure 5A, B**). Dominant frequencies of calcium spikes were determined by power spectrum analysis (**Figure 5C**). Consistent with our model, calcium spikes are fastest at the intermediate cross-linking level (**Figure 5D**). Next, we tested if the fastest calcium spikes result in a more frequent delivery of MT1-MMP vesicles to the invadopodial plasma membrane.

**Figure 5.**
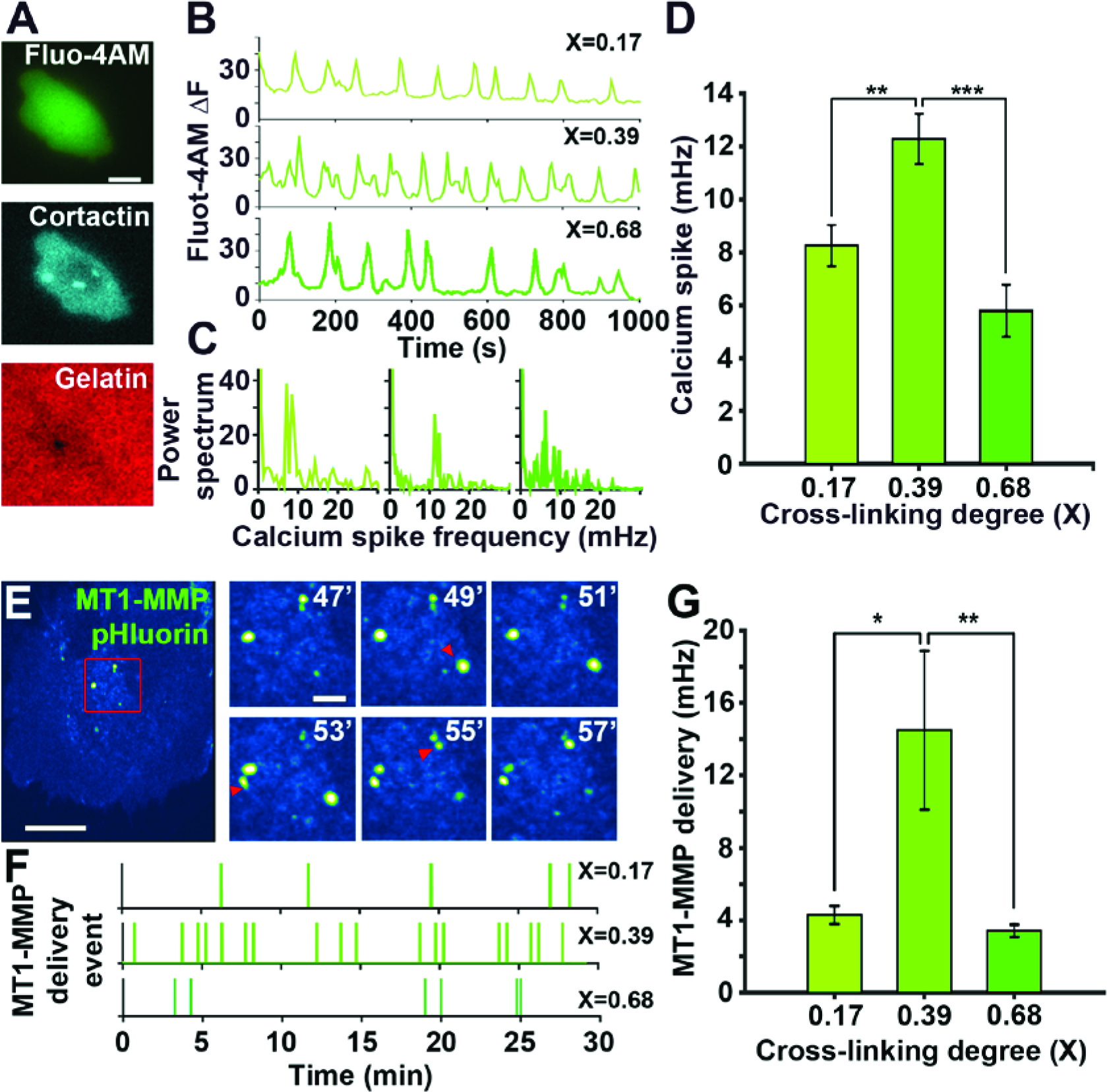
Frequencies of calcium spike and MT1-MMP vesicle delivery are biphasically regulated by ECM cross-linking. **A**. Representative fluorescence micrographs of Ceruleancortactin-MTLn3 (cyan, middle) cells (see **Movie S6**) labeled with Fluo-4-AM (green, top), plated on fluorescent gelatin (red, bottom). Scale bar is 10 microns; **B**. Representative calcium spikes recorded in cells at different gelatin cross-linking ratios; **C**. Power spectrum of calcium spikes in (**B**). **D**. Comparison of average spike frequencies at different gelatin cross-linking ratios. Scale bar 10 μm. Data includes >15 cells per experiment for 3 separate experiments per condition. Pairwise comparisons were done with 0.39 cross-linking ratio condition. **E.** Representative image of an HS-578T cell transfected with MT1-MMP pHluorin (see **Movie S7**). Right panel depicts dynamics of MT1-MMP containing vesicles delivery to the plasma membrane. Red arrows point at newly delivered vesicles. **F.** Representative record of MT1-MMP delivery event over 30 min in cells at different gelatin cross-linking ratios. **G.** Comparison of average frequency of MT1-MMP delivery at different gelatin cross-linking ratios. Data includes >15 cells per experiment for 3 separate experiments per condition. Pairwise comparisons were done with 0.39 cross-linking ratio condition. Errors are shown as SEM. P<0.01 **, P<0.001 ***.

### MT1-MMP containing vesicles delivery to the plasma membrane is biphasically regulated by ECM cross-linking

MT1-MMP is recycled by acidic late endosomes to the tip of mature invadopodia, where it is exocytosed to the neutral environment. We tested the MT1-MMP delivery rate using pH-sensitive MT1-MMPpHluorin. This fluorescent protein marks exocytic events of MT1-MMP, which appear as GFP-positive, blinking spots colocalized with the invadopodia (32). Exocytosis of MT1-MMPpHluorin vesicles was monitored at different cross-linking ratios (**Movie S6; Figure 5E, F**) to calculate the average vesicle delivery rate for each condition. At cross-linking ratio 0.39, MT1-MMP-vesicles were delivered more frequently compared to other cross-linking ratios (**Figure 5F, G**). Interestingly, frequencies of calcium spikes and MT1-MMP vesicle delivery operate in the similar range, suggesting a coordination between cycles of calcium spikes and vesicle delivery (31).

Collectively, our data suggests that frequencies of protrusive cycles in invadopodia as well as frequencies of calcium spikes and MT1-MMP delivery to the invadopodial plasma membrane change in concert and are controlled by ECM cross-linking.

### Switching between migration and Invadopodia states is regulated by β1-integrin activity

The coordinated relationship between ECM properties and invadopodia dynamics and function suggest a central role for the outside-in signaling provided by ECM-integrin interactions. As mentioned above, β1-integrin is localized to invadopodia and in its absence, invadopodia maturation and ECM-degrading function are disabled (7). We hypothesized that ECM-integrin β1 interactions are the master regulator of invadopodia dynamics and consequently, the length of time that cell will spend in Invadopodia state. We next tested this hypothesis in our model as well as experimentally.

In our model, the effect of cell-ECM interaction on invadopodia dynamics was examined by varying the adhesion constant (Ka) from 1 (**Figure 1E)** to 0.2 and 0.01 s^-1^ (**Figure 6A)**. A decrease in K_a_ reflects lower cell-ECM adhesion strength. While modifying ECM cross-linking alters adhesion strength externally by changing the number of ECM molecules proximal to the cell, adhesion strength can also be targeted by modifying the integrin ability to bind ECM. The model results suggest that a decrease in adhesion strength (K_a_ =0.2) will cause a decrease in the period spent in the Invadopodia state, simultaneously increasing the time spent in Migration state to the point where invadopodia is completely eliminated and the cell continuously migrates (K_a_=0.01).

**Figure 6.**
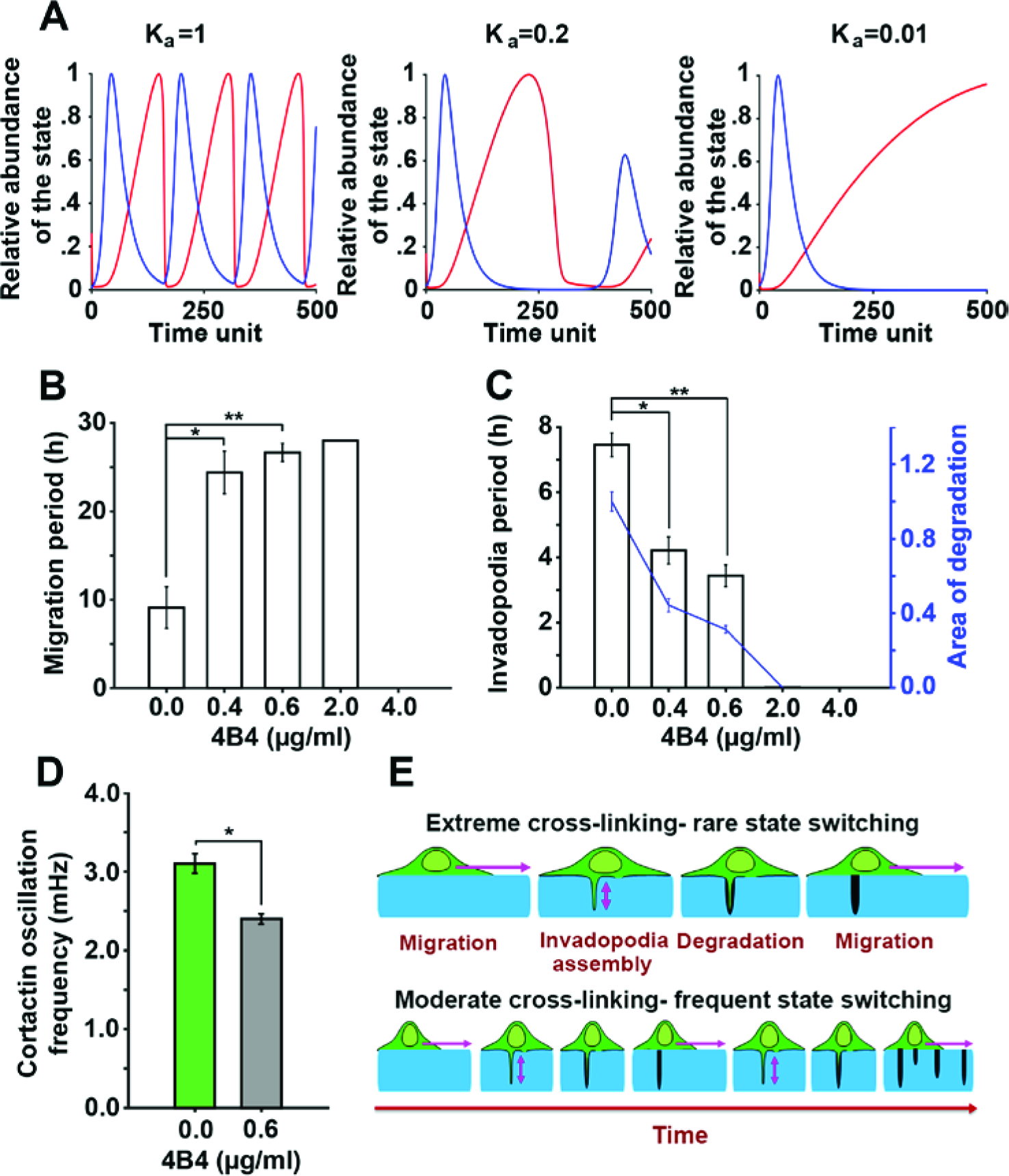
β1 integrin regulates balance between Migration/Invadopodia states. **A**. Simulation results of the cancer cell oscillation between invadopodia (blue) and migration (red) for decreasing values of K_a_, indicating elimination of Invadopodia state at K_a_ =0.01 (right panel); **B**. Change in time spent in Migration state (h) with increasing doses of β1 integrin blocker; **C.** Change in time spent in Invadopodia state (h) and invadopodia degradation with increasing doses of β1 integrin blocker 4B4. Blue line shows relative invadopodia degradation per cell (blue axis). Pairwise comparisons were done with 0 μg/ml 4B4 condition. **D.** Cortactin oscillation frequency measured in cells: treated with PBS (green bar) or cells treated with 0.6 μg/ml 4B4 β1 integrin inhibitor (grey bar). Experiments were done on gelatin layer with cross-linking degree of 0.36, in triplicate and errors are shown as SEM. * P<0.05; ** P<0.01. **E**. Summary of the conclusions. Levels of ECM cross-linking determines ECM-integrin interactions, which controls speed of invadopodia “digging” and ECM degradation. Faster invadopodia dynamics leads to more frequent switching between Invadopodia and Migration states and finally, to higher total ECM degradation.

To test the simulation results, β1-integrin ability to bind to the ECM was increasingly blocked using different doses of a monoclonal blocking antibody (4B4). The extent of invadopodia degradation and average lengths of both Invadopodia and Migration states, which occur on the timescale of hours, were measured at various doses of 4B4. Results show that with increasing concentration of 4B4, average time that a cell spends in Invadopodia state decreases while that of Migration state increases (**Figures 6B, C**). At 2.0 μg/ml of 4B4, ECM degradation is totally halted and cells migrate continuously. Higher concentrations of blocking antibody also block migration and cause cell detachment from the gelatin layer (≥4.0 μg/ml). Furthermore, we tested the effect of partial β1 integrin inhibition on the dynamics of invadopodia-related activities, such as cortactin oscillations, which occur on the timescale of minutes. Results show a significant decrease in the frequency of cortactin oscillations from 3.08 mHz, in control cells, to 2.39 mHz in cells with partial β1 integrin inhibition (**Figure 6D**). Such a decrease is reminiscent of the effect of extreme ECM cross-linking values (**Figure 4D**). Collectively, these results indicate that ECM-β1 integrin interactions are involved in regulating invadopodia-related dynamics on the timescale of minutes and in turn, frequency of switching between Invadopodia and Migration states, on the timescale of hours (**Figure 6E**).

## Discussion

Invadopodia assembly and function has been well studied as a measure of cancer cell invasiveness, but the relationship between invadopodia and cell translocation and the dynamics of these events were never directly addressed. Here, we demonstrate, for the first time, that cancer cells with invadopodia repeatedly oscillate between invadopodia and Migration states. Importantly, we show that the degree of ECM cross-linking controls the balance between the two states via the level of β1 integrin activity. Moreover, ECM cross-linking controls invadopodia dynamics and function, which involve protrusion-retraction cycles and calcium-dependent MT1-MMP delivery to plasma membrane.

The increase in ECM cross-linking has been previously demonstrated to increase the number of focal adhesions (22) and invadopodia (2, 14, 33). Further, stiffness of ECM has been reported to affect invadopodia numbers and activity (15). Finally, either an increase in ECM stiffness or mechanical stretching of ECM layer have been demonstrated to increase MMP expression (34, 35). Here, we show that the increase in ECM cross-linking affects invadopodia-related dynamics and their ECM degrading function. While the number of precursors plateaus with the increase in cross-linking, the number of mature invadopodia demonstrates a pronounced biphasic trend, suggesting that the cross-linking variations may be more important in later steps of invadopodia assembly, such as maturation and MT1-MMP delivery steps. Our data on MT1-MMP recycling confirms this hypothesis. Collectively, our data demonstrate that intermediate level of ECM cross-linking supports highest speeds of protrusive cycles, as well as the most frequent MT1-MMP delivery via Ca^2+^oscillations, while making invadopodia more stable, resulting in a peak of degradative activity. Furthermore, extent of ECM-β1-integrin interactions dictates the length of time that a cell can spend in Invadopodia state and the frequency of switching between migration and Invadopodia states.

Previous quantitative studies in both invadopodia, generated by cancer cells (13) and podosomes, generated by macrophages or dendritic cells (11, 36, 37), have shown an oscillatory behavior of the structure core, reflecting protrusion-retraction cycles. Intensity fluctuations in the core actin and cortactin content are a direct measure of the vertical movement of the protrusion tip” digging” into the ECM (11). Similar oscillations were seen in stiffness levels of the podosome structure itself, as measured by AFM (36). Lengths of protrusion-retraction cycles (i.e. core oscillations) reported in various cell types were 300-900 seconds (11, 13). Elimination of such cycles was seen with perturbations of actin core by inhibition of ROCK or myosin light chain kinase (11) or inhibition of cortactin phosphorylation (13). Our data suggests that the cell “sensing” of ECM cross-linking translates into frequency of protrusion-retraction cycles in a coordinated fashion. Thus, ECM cross-linking degree optimal for maximum degradation is in coordination with fastest protrusion-retraction cycles, as well as highest frequencies of calcium spikes and MT1-MMP delivery. Interestingly, coordination between calcium spikes and oscillations in cortical actin (as well as N-WASP and PI(4,5)P2) was recently demonstrated to be essential in vesicle secretion in leukemia cells (38, 39).

In breast carcinoma (7), as well as metastatic melanoma (40), β1-integrin is localized to invadopodia. β1-Integrin is also highly expressed in breast carcinoma compared to β3 and β5 and is necessary for invadopodium maturation, while β3 role focuses on general adhesions. Our model and results of blocking ECM-integrin β1 interactions suggest that the balance between migration and Invadopodia states can be altered via ECM-β1-integrin binding levels. Previous reports of dose-response to soluble factors such as EGF, demonstrated similar, biphasic trends of invadopodia numbers and/or chemotactic migration to increasing EGF concentrations (41–43). Taken together, these data suggest that conditions in the extracellular environment cannot be directly taken as having a positive or negative effect on motility or invadopodial functions. Cells continuously adapt their behavior to match the conditions encountered at extracellular environment, and this may include reverting to their invasive behaviors. It is possible that each stage of invadopodia assembly contains equilibrium points, and those invadopodia that do not reach stable conformation are continuously eliminated. Such a model is strengthened by a recent mathematical study on focal adhesion growth (44) suggesting that focal adhesions grow to the stable, equilibrium length. While the adhesions that did not reach equilibrium fully disassemble, those that surpass equilibrium size go through partial disassembly. Finally, stable focal adhesion size was found to be in direct relationship with ECM stiffness. In conclusion, our observations suggest the importance of considering non-linear relationships between cells and their immediate environments, which are further complicated by receptor transactivation and inherent heterogeneities in cell transcription levels.

Data presented can be summarized in the model where ECM-integrin β1 adhesion regulates dynamics of invadopodia-related processes occurring on the time scale of minutes (speed of protrusion-retraction cycles and ECM degradation). This in turn regulates dynamics of invadopodia-migration oscillations, which occurs on the time scale of hours. Our data demonstrates that faster protrusion-retraction cycles are in conjunction with faster MT1-MMP recycling and calcium spikes, leading to faster ECM degradation. Next, this leads to invadopodia disassembly and cell translocation to a place with intact ECM. In this model, integrin signaling regulates the speed and the efficiency of invadopodia short scale dynamics, therefore, regulating long scale dynamics of switches from cell Migration state to Invadopodia state (**Figure 6E**). Our mathematical modeling strengthens this view. By simulating a wide range of conditions in a generalizable, non-parametric manner, model predicts that changes in ECM-integrin binding and/or ECM cross-linking will induce changes in frequency of oscillations between Migration and Invadopodia states. Physically, this can be interpreted as requirement for non-overlapping levels of cell-ECM adherence in each of the states.

Our findings suggest the possibility of targeting invadopodia via ECM-modulation treatments. As invadopodia *in vivo* are necessary for intravasation (19) and extravasation (3) and hence, metastasis (19), understanding relationships between the ECM and invadopodia carries powerful implications for future chemotherapies. If invadopodia and their degradative activity can be destabilized by minute changes in ECM cross-linking, this might potentially turn off the cancer cell metastatic potential. *In vivo*, there are several enzymes catalyzing covalent cross-links of collagen and elastin in cancer. This includes LOX, PLOD and TGII enzymes, all of which were linked to increased cancer cell motility and metastasis (45–47). Additionally, ECM cross-linking can be induced by non-enzymatic glycation between sugars and proteins, resulting in advanced glycation products (AGEs), which were shown to increase invasion in cancer cells (48). Reduction of ECM cross-linking via inhibitors specifically designed for LOX, PLOD2 or TGII (42, 49, 50), as well as inhibitors of glycation such as flavonoids or aspirin (51) are likely to contribute towards invadopodia reduction. Our work suggests that such inhibitors can be valuable in reducing metastatic potential in neoadjuvant therapy.

## Methods

### Cell Culture

Mouse breast carcinoma line, Cerulean-cortactin-MTLn3, were a gift from Dr. Ved Sharma, Albert Einstein College of Medicine (Sharma et al. 2013). Cells were cultured in α-MEM (Gibco, Thermofisher Scientific, Waltham, MA) supplemented with 5 % FBS (Gemini Bio-Products, West Sacramento, CA), Penicillin/Streptomycin mixture (Gibco, Thermofisher Scientific, Waltham, MA) and 0.5 mg/ml G418 (Sigma-Aldrich). Cerulean-LifeAct-MDA-MB-231 and Cerulean-LifeAct-H2B-GFP cell lines were generated using mCerulean3-LifeAct-7 plasmid, selected with 0.5 mg/ml G418 over 4-week period and purified by FACS. mCerulean3-LifeAct-7 was a gift from Michael Davidson (Addgene plasmid #54721). Human cell lines MDA-MB-231 and Hs-578T (ATCC, Manassas, VA) cells were cultured in DMEM (Gibco, Thermofisher Scientific, Waltham, MA) supplemented with 10% FBS and 1% Penicillin/Streptomycin. For Hs-578T cells, 1 mM Pyruvate (Gibco, Thermofisher Scientific, Waltham, MA) and 0.01 mg/ml Bovine insulin (Sigma-Aldrich, St. Louis, MO) were added to the culture media.

### Cumulative degradation of gelatin layer

MTLn3, MDA-MB-231 and Hs578T cells were cultured on gelatin-coated plates (˜500 nm thickness) described previously (52). Briefly, acid-washed 35-mm MatTek (MatTek corporation, Ashland, MA) dishes were treated with 50 μg/ml Poly-L-Lysine (Gibco, Thermofisher Scientific, Waltham, MA) for 20 min followed by incubating with Alexa 488 dye-labeled gelatin (Sigma, St. Louis, MO) for 10 min. Plates were then washed with PBS (Gibco, Thermofisher Scientific, Waltham, MA) and crosslinked by glutaraldehyde (GTA, Sigma, St. Louis, MO) on ice for 15 min (from 0.0%-10.0 % v/v GTA in water), extensively rinsed with PBS, quenched with 5 mg/ml sodium borohydride (Sigma-Aldrich, St. Louis, MO), and sterilized with 70% ethanol (Decon Laboratories, King of Prussia, PA).

60,000 cells per dish were plated for 18 hours on 35 mm Mattek. Cells were fixed with 4% v/v paraformaldehyde (Electron Microscopy Sciences, Hatfield, PA) for 15 min and permeabilized with 0.1% Triton X-100 (Sigma-Aldrich, St. Louis, MO) for 10 min. Fixed cells were then blocked in 1% BSA (fraction V, Sigma-Aldrich, St. Louis, MO) and 1% FBS for 2 hours, incubated at 1:100 with anti-cortactin antibody (ab-33333, Abcam, Cambridge, UK) for 2 hours, followed by 2 hours incubation with Alexa-Fluor-633 (1:250) and Alexa-Fluor-546-Phalloidin (1:250) (Abcam, Cambridge, UK).

Imaging of the invadopodia precursors and cumulative degradation at 18 h were performed using an Olympus Fluoview 1200 confocal microscope (Olympus, Tokyo, Japan) with sequential imaging of 6 x 1 μm z-sections/stack.

Invadopodia degradation was assessed by quantifying the area of degradation per cell and the number of invadopodia precursors (colocalization of actin and cortactin) per cell at each condition in Fiji (53). For area of degradation, thresholding and particle size analysis were performed using Fiji. Experiments were done in triplicates, by imaging >10 fields of view, >100 cells in each sample.

### 3D FITC-DQ collagen I assay

100,000 MTLn3 cells were suspended in 100 μl of collagen mixture containing: 1.5 mg/ml collagen (Corning, Tewksbury, MA), 50 mM Tris-HCl, 2.5 mM CaCl2, 1 mM DTT (Sigma-Aldrich, St. Louis, MO), 0.15 mg/ml FITC-DQ collagen I (Molecular Probes, Thermofisher Scientific, Waltham, MA), and transglutaminase II (TGII; R&D Systems, Minneapolis, MN) at 1:50,000-1:800,000 dilution. Similarly to GTA, TGII crosslinks free amine groups of glutamine and lysine residues of collagen I (27). The mixture was pipetted in a 35-mm MatTek dish and incubated at 37°C for 30 min, forming a 650 μm-thick 3D cell culture. 2 ml of media (α-MEM/5% FBS) was pipetted into the plate. At 0 h and 42 h, 10 image stacks (20 x 5 μm) were collected. Olympus Fluoview FV1200 confocal microscope (Olympus, Tokyo, Japan) was used, with 488-nm laser excitation.

3D collagen degradation was assessed by measuring the area positive for DQ collagen at each condition. Briefly, images captured at 42h were thresholded and Particle Analysis was used in Fiji to quantify area positive for FITC-DQ collagen fluorescence. Measured values were subtracted from negative controls treated with GM6001 MMP inhibitor. All measurements were done in 10 fields of view per sample, with three biological repeats per condition.

### 3D collagen pore size measurement

3D cultures embedded in FITC-DQ collagen I and cross-linked to different degrees were imaged by reflection confocal microscopy. Images were thresholded with “Yen” algorithm in Fiji, filtered using LoG 3D plug-in and thresholded again with “Yen” to get the binary image of the pores. Finally, Particle Analysis was used to measure the size of the pores.

### Crosslinking degree measurements

Measurement of crosslinking degree followed previously published techniques (27, 54). Briefly, 0.2% w/v gelatin samples were mixed with 0.0%-5.0% v/v GTA and incubated on ice for 10 min. Crosslinking was quenched with 5 mg sodium borohydride for 15 minutes. For collagen, mixture of 1.5 mg/ml collagen I in PBS, 50 mM Tris-HCl, 2.5 mM CaCl2, 1mM DTT was crosslinked with 1:0.5K-1:800K TGII:collagen at 37°C for 30 minutes.

Uncrosslinked amino groups were neutralized by incubation with of 4% NaHCO and 0.5% trinitroben zenesulfonic acid (TNBS, Sigma, St. Louis, MO) for 2 h at 37°C in the dark. Finally, samples were incubated at 37°C for 1 h with 6 M HCL (Sigma, St. Louis, MO). The absorbance was measured at 345 nm using an Infinite M200 pro Tecan plate reader (Tecan, Männedorf, Switzerland).

### Live cell imaging

Live cell imaging was performed for invadopodia life time, cortactin oscillations, and calcium spike analysis. In brief, Cerulean-cortactin-MTLn3 cells were cultured on Alexa 488 or 546-gelatin coated MatTek dishes and incubated in culture conditions for 16 hours. For lifetime analysis and cortactin oscillations, media was changed to L15 with 5% FBS, 1:100 Oxyfluor (Oxyrase, Mansfield, OH), and 10 mM sodium lactate (Sigma-Aldrich, St. Louis, MO) as reactive oxygen scavengers. Dishes were placed in a 37 °C live cell imaging chamber and time lapse imaging was performed on widefield Olympus IX81 microscope (Olympus, Tokyo, Japan) equipped with LED lamp, Hamamatsu Orca 16-bit CCD (Hamamatsu, Hamamatsu city, Japan), automated z-drift compensation IX3-ZDC (Olympus, Tokyo, Japan), automated Prior stage (Prior Scientific, Rockland, MA) and environmental chamber. Olympus 60x 1.4 NA, Oil M Plan Apochromat objective was used.

### Measurement of invadopodia stability

Images were captured every 10 minutes for >16 hours (1.67 mHz). Invadopodia lifetimes were measured in Fiji and defined as the time between appearance and disappearance of cortactin punctae. Punctae present at the first or last frames were not taken into the account. Invadopodia stability ratio was defined as the ratio of the number of mature invadopodia (those with lifetime > 50 min (6)) to the total number of invadopodia.

### Cortactin oscillations in invadopodia core

Images were collected every 30 seconds (33.3 mHz) for 4 hours. Oscillations of cortactin fluorescence were measured only in mature invadopodia (present >50 min). For inhibiting F-actin polymerization, 4 μM Cytochalasin D (Sigma, St. Louis, MO) was added to the media prior to imaging. In Fiji we filtered individual invadopodia precursors with Laplacian of Gaussian (LOG 3d) (55) plugin and tracked the center of the invadopodium in all frames by SpotTracker (55). A custom written Fiji plugin was used to measure average signal inside the invadopodium as well as 4-8 circular areas inside the cytoplasm at each particular frame. In MATLAB, Fast Fourier Transform (FFT) was applied to the time-resolved cortactin signal in invadopodia and the surrounding cytoplasmic regions. High frequencies from the cytoplasm were removed and the filtered invadopodia signal was returned to the time domain by Inverse FFT. Finally, MATLAB autocorrelation function was used to measure the oscillation frequencies in invadopodia.

### Calcium Spikes

After cells were plated for 16 hours, cells were incubated for 15 minutes at room temperature in L15, 2.5 mM probenecid (Life Technologies, Frederick, MD), 2.5 μM Fluo4-AM (Life Technologies, Frederick, MD), and 0.02 % Pluronic (Life Technologies, Frederick, MD) (10). Cells were then washed twice with PBS and incubated in L15 with 5% FBS for 30 min at 37 C. Time lapse imaging was performed with 5 second intervals (200 mHz) for 1 hour. Cytoplasmic calcium spikes were measured by monitoring intracellular Fluo-4-AM signal over time (10). In Fiji, entire cells were used as regions of interest; signal was corrected for bleaching and background by subtracting the fluorescence in adjacent inactive cells. Measurements of inactive cells (˜70% of total, <10 spikes per hour) were not taken into account (10, 56). In MATLAB, power spectrum was applied to the time-resolved signal in the cytoplasm. Dominant frequencies at different conditions were further compared.

### Measurement of the MT1-MMP delivery event frequency

To visualize the delivery of MT1-MMP delivery event to the plasma membrane, Hs-578T cells were transiently transfected with MT1-MMPpHluorin 24 hours prior to imaging. Transfection was performed via an electroporation technique, in which 2 μg plasmid vector was mixed with 1x 10^6^cells and 50 μL nucleofection solution R (Lonza, Basel, Switzerland). Nucleofection was then performed on the mixture using program X-001 of a nucleofector device (Lonza, Basel, Switzerland). Transfected cells were then plated in a glass-bottom plate and 24 hours after plating, 30-minute time-lapse movies were captured from the cells using a 488 nm laser. A delivery event was defined as the moment that MT1-MMP flashes appear and the rate of delivery was measured as one over the average of the interval between successive flashes.

### β1 integrin blocking assay

The blocking of β1 integrin activity was done using 4B4 β1 integrin blocking antibody (Beckman Coulter) at doses of 0, 0.4, 0.6, and 2 μg/ml to cultures 2 hours after plating the cells. MDA-MB-231 cells were plated at a density of 60,000/x cm^2^ on Alexa 488 gelatin-coated dishes and time-lapse movies were recorded. Displacement of the cells over time was measured using Fiji plugin TrackMate. For area of degradation, thresholding and particle size analysis were performed using Fiji. Experiments were done in triplicates, by imaging >10 fields of view, >100 cells in each sample.

### Statistical Analysis

One way analysis of variance (ANOVA) statistical analysis with Tukey’s multiple comparison post-tests was performed to compare the 2D and 3D invadopodia degradation data. Two-tailed student's t-test was performed for statistical analysis as indicated and statistical significance was defined as * P < 0.05; ** P < 0.01 and *** P < 0.001. Data are shown as means ± SEM.

## Author Contributions

AB, KE and BG have designed, analyzed and interpreted all experiments and mathematical model and wrote the manuscript. KE has performed all experiments.

## Acknowledgements

We would like to thank Dr. Ved Sharma and the lab of Dr. John Condeelis for the gift of Cerulean-cortactin-MTLn3 cell line. We would like to also thank Dr. Philippe Chavrier for the gift of MT1-MMPpHluorin plasmid. We thank Dr. Edna Cukierman. Dr. Evangelia Bellas and Dr. Erica Golemis for help with editing this manuscript.

Authors declare no competing interests. This work was funded by NIH 5K99CA172360 and Concern Foundation grant to BG and NIH R01CA164468 and R01 DA033788 to AB.

## References

1. Monsky, W.L., C. Lin, A. Aoyama, T. Kelly, S.K. Akiyama, S.C. Mueller, and W.-T. Chen. 1994. A Potential Marker Protease of Invasiveness, Separase, Is Localized on Invadopodia of Human Malignant Melanoma Cells. Cancer Res. 54: 5702–5710.

2. Gligorijevic, B., A. Bergman, and J. Condeelis. 2014. Multiparametric Classification Links Tumor Microenvironments with Tumor Cell Phenotype. PLoS Biol. 12: e1001995.

3. Leong, H.S., A.E. Robertson, K. Stoletov, S.J. Leith, C.A. Chin, A.E. Chien, M.N. Hague, A. Ablack, K. Carmine-Simmen, V.A. McPherson, C.O. Postenka, E.A. Turley, S.A. Courtneidge, A.F. Chambers, and J.D. Lewis. 2014. Invadopodia Are Required for Cancer Cell Extravasation and Are a Therapeutic Target for Metastasis. Cell Rep. 8: 1558–1570.

4. Oser, M., H. Yamaguchi, C.C. Mader, J.J. Bravo-Cordero, M. Arias, X. Chen, V. DesMarais, J. Van Rheenen, A.J. Koleske, and J. Condeelis. 2009. Cortactin regulates cofilin and N-WASp activities to control the stages of invadopodium assembly and maturation. J. Cell Biol. 186: 571–587.

5. Artym, V. V., Y. Zhang, F. Seillier-Moiseiwitsch, K.M. Yamada, and S.C. Mueller. 2006. Dynamic interactions of cortactin and membrane type 1 matrix metalloproteinase at invadopodia: Defining the stages of invadopodia formation and function. Cancer Res. 66: 3034–3043.

6. Beaty, B.T., and J. Condeelis. 2014. Digging a little deeper: The stages of invadopodium formation and maturation. Eur. J. Cell Biol. 93: 438–444.

7. Beaty, B.T., V.P. Sharma, J.J. Bravo-Cordero, M.A. Simpson, R.J. Eddy, A.J. Koleske, and J. Condeelis. 2013. β1 integrin regulates Arg to promote invadopodial maturation and matrix degradation. Mol. Biol. Cell. 24: 1661–75, S1-11.

8. Beaty, B.T., Y. Wang, J.J. Bravo-Cordero, V.P. Sharma, V. Miskolci, L. Hodgson, and J. Condeelis. 2014. Talin regulates moesin-NHE-1 recruitment to invadopodia and promotes mammary tumor metastasis. J. Cell Biol. 205: 737–751.

9. Monteiro, P., C. Rossé, A. Castro-Castro, M. Irondelle, E. Lagoutte, P. Paul-Gilloteaux, C. Desnos, E. Formstecher, F. Darchen, D. Perrais, A. Gautreau, M. Hertzog, and P. Chavrier. 2013. Endosomal WASH and exocyst complexes control exocytosis of MT1-MMP at invadopodia. J. Cell Biol. 203: 1063–1079.

10. Sun, J., F. Lu, H. He, J. Shen, J. Messina, R. Mathew, D. Wang, A.A. Sarnaik, W.C. Chang, M. Kim, H. Cheng, and S. Yang. 2014. STIM1- and Orai1-mediated Ca2+ oscillation orchestrates invadopodium formation and melanoma invasion. J. Cell Biol. 207: 535–548.

11. van den Dries, K., M.B.M. Meddens, S. de Keijzer, S.C. Shekhar, V. Subramaniam, C.G. Figdor, and A. Cambi. 2013. Interplay between myosin IIA-mediated contractility and actin network integrity orchestrates podosome composition and oscillations. Nat. Commun. 4: 1412.

12. Steffen, A., G. Le Dez, R. Poincloux, C. Recchi, P. Nassoy, K. Rottner, T. Galli, and P. Chavrier. 2008. MT1-MMP-Dependent Invasion Is Regulated by TI-VAMP/VAMP7. Curr. Biol. 18: 926–931.

13. Magalhaes, M.A.O., D.R. Larson, C.C. Mader, J.J. Bravo-Cordero, H. Gil-Henn, M. Oser, X. Chen, A.J. Koleske, and J. Condeelis. 2011. Cortactin phosphorylation regulates cell invasion through a pH-dependent pathway. J. Cell Biol. 195: 903–920.

14. Enderling, H., N.R. Alexander, E.S. Clark, K.M. Branch, L. Estrada, C. Crooke, J. Jourquin, N. Lobdell, M.H. Zaman, S.A. Guelcher, A.R.A. Anderson, and A.M. Weaver. 2008. Dependence of invadopodia function on collagen fiber spacing and cross-linking: computational modeling and experimental evidence. Biophys. J. 95: 2203–2218.

15. Parekh, A., N.S. Ruppender, K.M. Branch, M.K. Sewell-Loftin, J. Lin, P.D. Boyer, O.E. Candiello, W.D. Merryman, S.A. Guelcher, and A.M. Weaver. 2011. Sensing and modulation of invadopodia across a wide range of rigidities. Biophys. J. 100: 573–582.

16. Alexander, N.R., K.M. Branch, A. Parekh, E.S. Clark, I.C. Iwueke, S.A. Guelcher, and A.M. Weaver. 2008. Extracellular Matrix Rigidity Promotes Invadopodia Activity. Curr. Biol. 18: 1295–1299.

17. Yu, X., and L.M. Machesky. 2012. Cells assemble invadopodia-like structures and invade into matrigel in a matrix metalloprotease dependent manner in the circular invasion assay. PLoS One. 7: e30605.

18. Tolde, O., D. Rösel, P. Veselý, P. Folk, and J. Brábek. 2010. The structure of invadopodia in a complex 3D environment. Eur. J. Cell Biol. 89: 674–680.

19. Gligorijevic, B., J. Wyckoff, H. Yamaguchi, Y. Wang, E.T. Roussos, and J. Condeelis. 2012. N-WASP-mediated invadopodium formation is involved in intravasation and lung metastasis of mammary tumors. J. Cell Sci. 125: 724–734.

20. Wolf, K., M. te Lindert, M. Krause, S. Alexander, J. te Riet, A.L. Willis, R.M. Hoffman, C.G. Figdor, S.J. Weiss, and P. Friedl. 2013. Physical limits of cell migration: Control by ECM space and nuclear deformation and tuning by proteolysis and traction force. J. Cell Biol. 201: 1069–1084.

21. Zaman, M.H., L.M. Trapani, A.L. Sieminski, D. MacKellar, H. Gong, R.D. Kamm, A. Wells, D.A. Lauffenburger, and P. Matsudaira. 2006. Migration of tumor cells in 3D matrices is governed by matrix stiffness along with cell-matrix adhesion and proteolysis. Proc. Natl. Acad. Sci. 103: 10889–10894.

22. Levental, K.R., H. Yu, L. Kass, J.N. Lakins, M. Egeblad, J.T. Erler, S.F.T. Fong, K. Csiszar, A. Giaccia, W. Weninger, M. Yamauchi, D.L. Gasser, and V.M. Weaver. 2009. Matrix Crosslinking Forces Tumor Progression by Enhancing Integrin Signaling. Cell. 139: 891–906.

23. Ehrbar, M., A. Sala, P. Lienemann, A. Ranga, K. Mosiewicz, A. Bittermann, S.C. Rizzi, F.E. Weber, and M.P. Lutolf. 2011. Elucidating the role of matrix stiffness in 3D cell migration and remodeling. Biophys. J. 100: 284–293.

24. Hoshino, D., N. Koshikawa, T. Suzuki, V. Quaranta, A.M. Weaver, M. Seiki, and K. Ichikawa. 2012. Establishment and validation of computational model for MT1-MMP dependent ECM degradation and intervention strategies. PLoS Comput. Biol. 8: e1002479.

25. Palecek, S.P., J.C. Loftus, M.H. Ginsberg, D.A. Lauffenburger, and A.F. Horwitz. 1997. Integrin – ligand binding properties govern cell migration speed through cell – substratum adhesiveness. Nature. 388: 537–540.

26. Li, A., J.C. Dawson, M. Forero-Vargas, H.J. Spence, X. Yu, I. Konig, K. Anderson, and L.M. Machesky. 2010. The Actin-Bundling Protein Fascin Stabilizes Actin in Invadopodia and Potentiates Protrusive Invasion. Curr. Biol. 20: 339–345.

27. Orban, J.M., L.B. Wilson, J. a Kofroth, M.S. El-Kurdi, T.M. Maul, and D. a Vorp. 2004. Crosslinking of collagen gels by transglutaminase. J. Biomed. Mater. Res. A. 68: 756–762.

28. Lyford, G.L., P.R. Strege, A. Shepard, Y. Ou, L. Ermilov, S.M. Miller, S.J. Gibbons, J.L. Rae, J.H. Szurszewski, G. Farrugia, L. Greg, P.R. Strege, A. Shepard, L. Ermilov, S.M. Miller, J. Simon, J.L. Rae, and J.H. Szurszewski. 2002. a 1C ( Ca V 1 . 2) L-type calcium channel mediates mechanosensitive calcium regulation. 55905: 1001–1008.

29. Farrugia, G., A.N. Holm, A. Rich, M.G. Sarr, J.H. Szurszewski, and J.L. Rae. 1999. A mechanosensitive calcium channel in human intestinal smooth muscle cells. Gastroenterology. 117: 900–905.

30. Calabrese, B., I. V Tabarean, P. Juranka, and C.E. Morris. 2002. Mechanosensitivity of N-type calcium channel currents. Biophys J. 83: 2560–2574.

31. Godbout, C., L. Follonier Castella, E.A. Smith, N. Talele, M.L. Chow, A. Garonna, and B. Hinz. 2013. The Mechanical Environment Modulates Intracellular Calcium Oscillation Activities of Myofibroblasts. PLoS One. 8: e64560.

32. Lizarraga, F., R. Poincloux, M. Romao, G. Montagnac, G. Le Dez, I. Bonne, G. Rigaill, G. Raposo, and P. Chavrier. 2009. Diaphanous-related formins are required for invadopodia formation and invasion of breast tumor cells. Cancer Res. 69: 2792–2800.

33. Lauzier, A., M. Charbonneau, M. Paquette, K. Harper, and C.M. Dubois. 2012. Transglutaminase 2 cross-linking activity is linked to invadopodia formation and cartilage breakdown in arthritis. Arthritis Res. Ther. 14: R159.

34. Milkiewicz, M., F. Mohammadzadeh, E. Ispanovic, E. Gee, and T.L. Haas. 2007. Static strain stimulates expression of matrix metalloproteinase-2 and VEGF in microvascular endothelium via JNK- and ERK-dependent pathways. J. Cell. Biochem. 100: 750–761.

35. Seo, K.W., S.J. Lee, Y.H. Kim, J.U. Bae, S.Y. Park, S.S. Bae, and C. Da Kim. 2013. Mechanical Stretch Increases MMP-2 Production in Vascular Smooth Muscle Cells via Activation of PDGFR-ß/Akt Signaling Pathway. PLoS One. 8: e70437.

36. Labernadie, A., C. Thibault, C. Vieu, I. Maridonneau-Parini, and G.M. Charrière. 2010. Dynamics of podosome stiffness revealed by atomic force microscopy. Proc. Natl. Acad. Sci. U. S. A. 107: 21016–21021.

37. Labernadie, A., A. Bouissou, P. Delobelle, S. Balor, R. Voituriez, A. Proag, I. Fourquaux, C. Thibault, C. Vieu, R. Poincloux, G.M. Charrière, and I. Maridonneau-Parini. 2014. Protrusion force microscopy reveals oscillatory force generation and mechanosensing activity of human macrophage podosomes. Nat. Commun. 5: 5343.

38. Smedler, E., and P. Uhlén. 2014. Frequency decoding of calcium oscillations. Biochim. Biophys. Acta - Gen. Subj. 1840: 964–969.

39. Wollman, R., and T. Meyer. 2012. Coordinated oscillations in cortical actin and Ca2+ correlate with cycles of vesicle secretion. Nat. Cell Biol. 14: 1261–9.

40. Mueller, S.C., G. Ghersi, S.K. Akiyama, Q.X.A. Sang, L. Howard, M. Pineiro-Sanchez, H. Nakahara, Y. Yeh, and W.T. Chen. 1999. A novel protease-docking function of integrin at invadopodia. J. Biol. Chem. 274: 24947–24952.

41. Philippar, U., E.T. Roussos, M. Oser, H. Yamaguchi, H. Do Kim, S. Giampieri, Y. Wang, S. Goswami, J.B. Wyckoff, D.A. Lauffenburger, E. Sahai, J.S. Condeelis, and F.B. Gertler. 2008. A Mena Invasion Isoform Potentiates EGF-Induced Carcinoma Cell Invasion and Metastasis. Dev. Cell. 15: 813–828.

42. Zhou, Z.N., V.P. Sharma, B.T. Beaty, M. Roh-Johnson, E.A. Peterson, N. Van Rooijen, P.A. Kenny, H.S. Wiley, J.S. Condeelis, and J.E. Segall. 2014. Autocrine HBEGF expression promotes breast cancer intravasation, metastasis and macrophage-independent invasion in vivo. Oncogene. 33: 3784–93.

43. Tönisen, F., L. Perrin, B. Bayarmagnai, K. van den Dries, A. Cambi, and B. Gligorijevic. 2017. EP4 receptor promotes invadopodia and invasion in human breast cancer. Eur. J. Cell Biol. 96: 218–226.

44. Cao, X., Y. Lin, T.P. Driscoll, J. Franco-Barraza, E. Cukierman, R.L. Mauck, and V.B. Shenoy. 2015. A Chemomechanical Model of Matrix and Nuclear Rigidity Regulation of Focal Adhesion Size. Biophys. J. 109: 1807–1817.

45. Mangala, L.S., B. Arun, A. a Sahin, and K. Mehta. 2005. Tissue transglutaminase-induced alterations in extracellular matrix inhibit tumor invasion. Mol. Cancer. 4: 33.

46. Joice, R., S.K. Nilsson, J. Montgomery, S. Dankwa, B. Morahan, K.B. Seydel, L. Bertuccini, P. Alano, C. Kim, M.T. Duraisingh, T.E. Taylor, and D.A. Milner. 2014. Procollagen Lysyl Hydroxylase 2 is Essential for Hypoxia- Induced Breast Cancer Metastasis. 6: 1–16.

47. El-Haibi, C.P., G.W. Bell, J. Zhang, A.Y. Collmann, D. Wood, C.M. Scherber, E. Csizmadia, O. Mariani, C. Zhu, A. Campagne, M. Toner, S.N. Bhatia, D. Irimia, A. Vincent-Salomon, and A.E. Karnoub. 2012. Critical role for lysyl oxidase in mesenchymal stem cell-driven breast cancer malignancy. Proc. Natl. Acad. Sci. 109: 17460–17465.

48. Sharaf, H., S. Matou-Nasri, Q. Wang, Z. Rabhan, H. Al-Eidi, A. Al Abdulrahman, and N. Ahmed. 2015. Advanced glycation endproducts increase proliferation, migration and invasion of the breast cancer cell line MDA-MB-231. Biochim. Biophys. Acta - Mol. Basis Dis. 1852: 429–441.

49. Eisinger-Mathason, T.S.K., M. Zhang, Q. Qiu, N. Skuli, M.S. Nakazawa, T. Karakasheva, V. Mucaj, J.E.S. Shay, L. Stangenberg, N. Sadri, E. Puré, S.S. Yoon, D.G. Kirsch, and M.C. Simon. 2013. Hypoxia-dependent modification of collagen networks promotes sarcoma metastasis. Cancer Discov. 3: 1190–1205.

50. Jeitner, T.M., E.J. Delikatny, J. Ahlqvist, H. Capper, and A.J.L. Cooper. 2005. Mechanism for the inhibition of transglutaminase 2 by cystamine. Biochem. Pharmacol. 69: 961–970.

51. Peng, X., J. Ma, F. Chen, and M. Wang. 2011. Naturally occurring inhibitors against the formation of advanced glycation end-products. Food Funct. 2: 289.

52. Sharma, V.P., D. Entenberg, and J. Condeelis. 2013. High-resolution live-cell imaging and time-lapse microscopy of invadopodium dynamics and tracking analysis. Methods Mol. Biol. 1046: 343–57.

53. Schindelin, J., I. Arganda-Carreras, E. Frise, V. Kaynig, M. Longair, T. Pietzsch, S. Preibisch, C. Rueden, S. Saalfeld, B. Schmid, J.-Y. Tinevez, D.J. White, V. Hartenstein, K. Eliceiri, P. Tomancak, and A. Cardona. 2012. Fiji: an open-source platform for biological-image analysis. Nat. Methods. 9: 676–682.

54. Grover, C.N., J.H. Gwynne, N. Pugh, S. Hamaia, R.W. Farndale, S.M. Best, and R.E. Cameron. 2012. Crosslinking and composition influence the surface properties, mechanical stiffness and cell reactivity of collagen-based films. Acta Biomater. 8: 3080–3090.

55. Sage, D., F.R. Neumann, F. Hediger, S.M. Gasser, and M. Unser. 2005. Automatic tracking of individual fluorescence particles: Application to the study of chromosome dynamics. IEEE Trans. Image Process. 14: 1372–1383.

56. Hamadi, A., G. Giannone, K. Takeda, and P. Rondé. 2014. Glutamate involvement in calcium-dependent migration of astrocytoma cells. Cancer Cell Int. 14: 42.

57. Lauffenburger, D. a, and a L. Horwitz. 1996. Cell migration: A physically integrated process. Cell. 84: 359–369.

58. DiMilla, P.A., K. Barbee, and D.A. Lauffenburger. 1991. Mathematical model for the effects of adhesion and mechanics on cell migration speed. Biophys. J. 60: 15–37.

59. Ohuchi, E., K. Imai, Y. Fujii, H. Sato, M. Seiki, and Y. Okada. 1997. Membrane-Type metalloproteinase digests extracellular matrix macromolecules including interstitial collagens. Matrix Biol. 16: 76–77.

